# ACE2 expression in rat brain: implications for COVID-19 associated neurological manifestations

**DOI:** 10.1101/2021.05.01.442293

**Authors:** Vito S Hernández, Mario A Zetter, Enrique C. Guerra, Ileana Hernández-Araiza, Nikita Karuzin, Oscar R. Hernández-Pérez, Lee E Eiden, Limei Zhang

## Abstract

We examined cell type-specific expression and distribution of rat brain angiotensin converting enzyme 2 (ACE2), the receptor for SARS-CoV-2, in rodent brain. ACE2 is ubiquitously present in brain vasculature, with the highest density of ACE2 expressing capillaries found in the olfactory bulb, the hypothalamic paraventricular, supraoptic and mammillary nuclei, the midbrain substantia nigra and ventral tegmental area, and the hindbrain pontine nucleus, pre-Bötzinger complex, and nucleus of *tractus solitarius*. ACE2 was expressed in astrocytes and astrocytic foot processes, pericytes and endothelial cells, key components of the blood-brain-barrier. We found discrete neuronal groups immunopositive for ACE2 in brainstem respiratory rhythm generating centers including the pontine nucleus, the parafascicular/retrotrapezoid nucleus, the parabrachial nucleus, the Bötzinger and pre-Bötzinger complex and the nucleus of tractus solitarius; in arousal-related pontine reticular nucleus and in gigantocellular reticular nuclei; in brainstem aminergic nuclei, including substantia nigra, ventral tegmental area, dorsal raphe, and locus coeruleus; in the epithalamic habenula, hypothalamic paraventricular and suprammamillary nuclei; and in the hippocampus. Identification of ACE2-expressing neurons in rat brain within well-established functional circuits facilitates prediction of possible neurological manifestations of brain ACE2 dysregulation during and after COVID-19 infection.

**Highlights:**

- ACE2 is present in astrocytes, pericytes, and endothelia of the blood brain barrier.
- Neuronal ACE2 expression is shown in discrete nuclei through the brain.
- Brainstem breathing, arousal-related, hypothalamic and limbic nuclei express ACE2.
- ACE2 is expressed in circuits potentially involved in COVID-19 pathophysiology.

## Introduction

The angiotensin converting enzyme ACE2 (EC 3.4.15.1), is a metalloproteinase that was discovered in 2000 by two independent groups (Donoghue, Hsieh et al. 2000, Tipnis, Hooper et al. 2000). Since then, it has been characterized as a counterregulatory component for the classical renin-angiotensin-aldosterone system (RAAS), responsible for cleaving angiotensins I and II to peptides (angiotensin (1-9) and (1-7)) whose effects oppose the vasoconstrictor/proinflammatory actions of angiotensins generated by angiotensin converting enzyme (ACE) (Rice, Thomas et al. 2004) (Zhang, Zetter et al. 2021). Moreover, the ability of ACE2 to hydrolyze other peptides such as apelin, kinins, (des-Arg9)-bradykinin, neurotensin and dynorphin A-(1-13) (Vickers, Hales et al. 2002) provide additional complexity to the role(s) of ACE2 in RAAS counter-regulation, and association with various disease pathologies.

SARS-CoV-2, the pathogen of the current COVID-19 pandemic, is associated with the RAAS (Zhang, Zetter et al. 2021). The virus uses ACE2 as a receptor to invade cells by binding to it via the viral trimeric spike protein (Yan, Zhang et al. 2020). The spike protein is primed by the serine protease TMPRSS2, triggering the fusion of viral and cellular membranes, and internalization of both virus and receptor in the first step of cellular infection (Zhang, Penninger et al. 2020).

The acute and chronic neurological manifestations of COVID-19, such as headache, dizziness, loss or disruption of the sense of smell (anosmia/dysosmia), taste (ageusia/dysgeusia), loss of muscular coordination (ataxia), loss of autonomic respiratory control, lethargy, depression and anxiety (Kabbani and Olds 2020, Mao, Jin et al. 2020, Satarker and Nampoothiri 2020, Haidar, Jourdi et al. 2021, Nagu, Parashar et al. 2021, Stefano, Ptacek et al. 2021), as well as its potential contribution to long-term adverse cerebrovascular, neuropsychiatric, and neurodegenerative pathologies (Cohen, Eichel et al. 2020, Ding, Shults et al. 2020, Nagu, Parashar et al. 2021, Rahman, Islam et al. 2021) (Sashindranath and Nandurkar 2021) are still poorly understood. Thus, there is an urgent need to determine brain ACE2 expression at the cellular and regional levels and to analyze its potential participation in defined functional circuits associated with the neurological manifestations of COVID-19.

Here, we examined cell type-specific expression, and regional distribution of ACE2 in rodent brain. We used immunohistochemistry to characterize the pattern of ACE2 expression in vascular, glial, and neuronal elements, participants of several well-defined circuits for respiratory rhythm, arousal, reward, homeostasis, learning and memory. We analyze and discuss the possible consequences of the disruption of these circuits in contributing to COVID-19 CNS disease.

## Material and methods

### Animals and brain section preparation

Four male Wistar rats from the local animal breeding facility were used in the present study. Animals were housed with two other littermates, with food and water *ad libitum*, temperature maintained between 20°C and 25°C, artificial illumination established to light-on at 8:00 hrs. and light-off at 20 hrs. and bedding changed three times per week. For immunohistochemistry, rats were deeply anesthetized with pentobarbital 63 mg/kg; when eyelid reflex was abolished, the animals were perfused transcardially with 0.9% NaCl solution followed by 0.1 M phosphate buffer (PB, pH 7.4) containing 4% paraformaldehyde and 15% v/v of a saturated picric acid solution. The brains were quickly removed and thoroughly washed with PB. Sagittal 70 μm sections through the whole mediolateral span were obtained with a vibratome (Leica VT1000S, Germany). All animals were handled according to the guidelines and requirements of the National Institutes of Health Guide for the Care and Use of Laboratory Animals (8th edition), and the Mexican Official Norm for Use, Care and Reproduction of Laboratory Animals (NOM-062-ZOO-1999). Experimental protocols were reviewed and approved by the local Research and Ethics Committee (CIEFM-062-2016).

### Immunohistochemistry

For immunoperoxidase staining for ACE2, sagittal serial sections (at intervals of 140 μm) were selected from two rats. Sections were first blocked against unspecific labeling through incubation in 10% normal donkey serum (NDS) in 0.05 M Trizma buffer, with 0.9% NaCl and 0.3% Triton X-100 (TBST), at room temperature (RT) for two hours. Primary antibody rabbit anti-ACE2 (ab15348, 1:1000, Abcam, MA, USA, see also table 1 for antibody information) was used, diluted in TBST + 1% NDS during 48h at 4°C with gentle shaking. After being washed with TBST, sections were incubated with a peroxidase-conjugated donkey anti-rabbit IgG (Jackson Immunoresearch, 711-035-152, PA, USA) for 2 hrs. at RT. Sections were washed with 0.1 M PB and immunoreaction was chromogenically developed using 3,3’-diaminobenzidine (Sigma-Aldrich, D5637, MO, USA) and H_2_O_2_ as substrates. Negative control sections were processed without adding the primary antibody. Finally, slices were mounted on glass slides, dehydrated with ethanol, transferred to xylene and cover-slipped using Permount mounting medium. (Fischer Chemical, SP15-100, MA, USA).

**Table 1.**
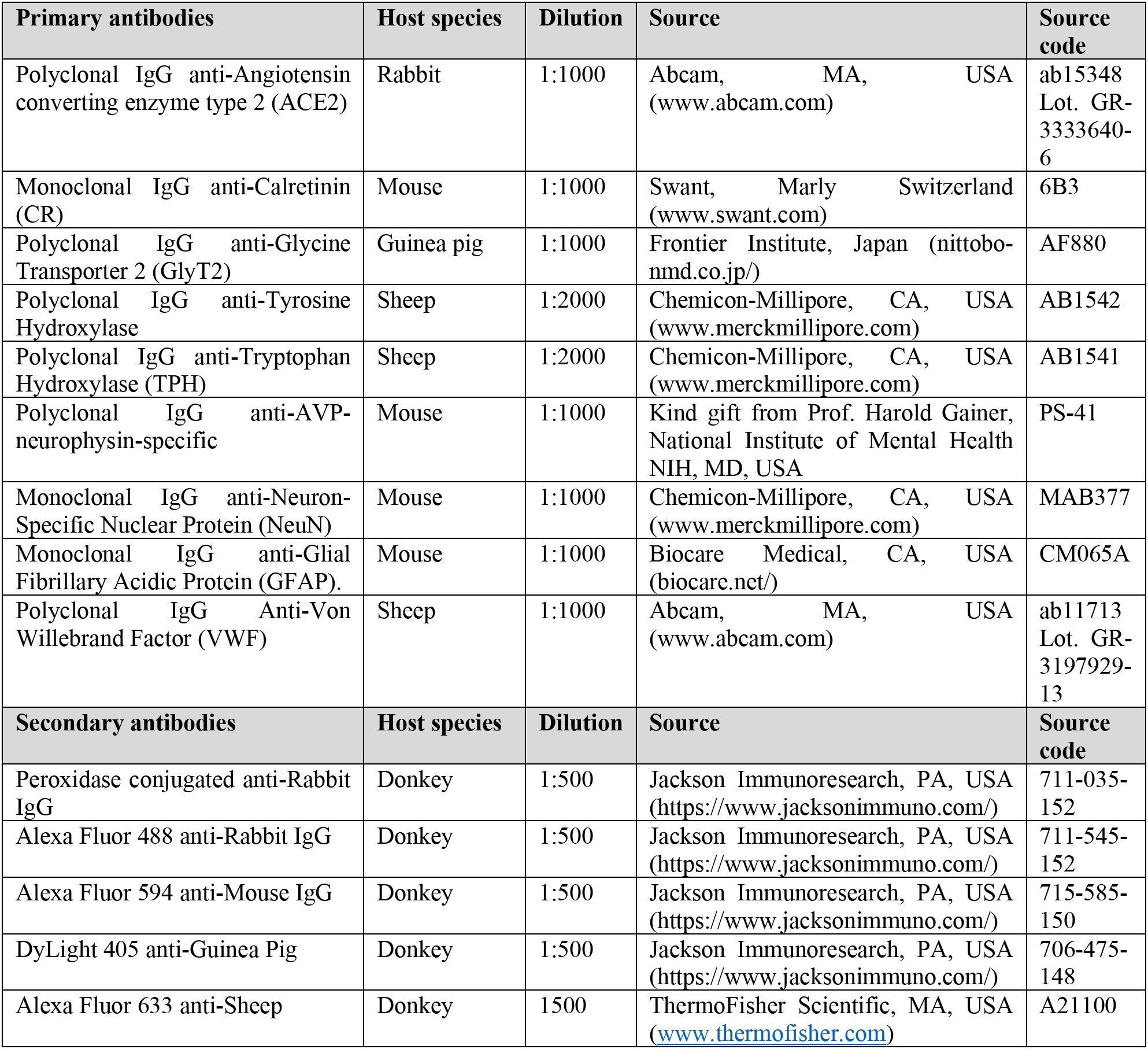
Primary and secondary antibodies

For multi-channel immunofluorescence staining, sections adjacent to those processed for chromogenic reaction were selected and incubated with 10 % normal donkey serum in TBST for two hr. at RT. After this blocking step, sections were incubated during 48hr at 4°C with cocktails of primary antibodies against ACE2 and combinations of the following markers: calretinin (CR, a calcium binding protein with a high expression in several nuclei of the brainstem and in the olfactory bulb); the glycine transporter 2 (GlyT2, a marker of glycinergic neurons, highly expressed in the pontomedullary region of the brain); tyrosine hydroxylase (TH, a marker for catecholaminergic neurons), tryptophan hydroxylase (TPH, a marker for serotoninergic neurons); AVP-neurophysin (a marker for vasopressinergic neurons); neuron-specific nuclear protein (NeuN, a pan-neuronal marker); Von Willebrand Factor (VWF, a marker for endothelial cells) and glial fibrillary acidic protein (GFAP, a marker for astrocytes). See Table 1 for details about the antibodies used in this study. After primary antibody incubation, the sections were washed three times in TBST and incubated with the corresponding secondary antibodies (see table 1). Finally, sections were washed thoroughly in TBS and mounted on glass slides with Vectashield antifade mounting medium (Vector, H1000, CA, USA). The DAB developed slices were observed with an Olympus CX31 light microscope. Panoramic reconstructions of chromogenic developed sections were obtained with a Zeiss AxioZoom V.16 microscope. Fluorescent images were obtained using a Zeiss confocal LSM 880.

### Quantification of ACE2 expression by optical intensity measurement in immunofluorescence photomicrographs

Using the whole-brain digitally scanned images of the chromogenically developed slides, we identified regions of high vascularization. A confocal fluorescence image was obtained from similar regions in the adjacent sections that were processed for immunofluorescence. Laser intensity, Airy disk diameter, detector gain, offset and exposure time parameters were fixed between photomicrographs. Images were obtained in an 8bit format, thus the possible values for each pixel ranged from “0” (no light) to “255” (maximum possible value). To calculate the optical intensity (O.I.) from the light emitted at a determined region of interest (ROI), the average O.I. of the pixels in a square 2500 μm^2^area within the ROI was calculated in the program ZEN lite 3.1 (Carl Zeiss Microscopy GmbH, Germany). The average O.I. value of the background was calculated from five regions where there was no tissue, thus only obtaining the value from the glass slide and mounting medium, this background value was subtracted from the value obtained in the ROIs. Finally, O.I. values were standardized, assigning a value of 10 relative fluorescence intensity units to the main olfactory bulb (the region with the highest measured O.I.).

## Results

### General expression pattern of ACE2

The initial evaluation of ACE2 immunoreactivity through the rat brain was performed in peroxidase-DAB developed slices (figure 1). High expression of ACE2 was observed in capillaries throughout the brain, with the highest expression in the glomerular layer of the main olfactory bulb (MOBgl). Other regions with a high density of ACE2 expressing vessels were the supraoptic (SON), paraventricular (PVN) and mammillary (MM) nuclei of the hypothalamus; the lateral habenula (LHb) in the epithalamus; the *substantia nigra* (SN) and ventral tegmental area (VTA) in the mesencephalon; and the nucleus of tractus solitarius (NTS), gigantocellular reticular nucleus (GRN), pontine nucleus (PN), pontine reticular nucleus (PRN), pre-Bötzinger complex (pre-BötC) and Bötzinger complex within the pontomedullary region. ACE2 expression was quantified by measurement of the relative fluorescence intensity (*RFI*) in photomicrographs obtained from adjacent slices from the regions shown and processed for ACE immunofluorescence (Table 2).

**Fig. 1.**
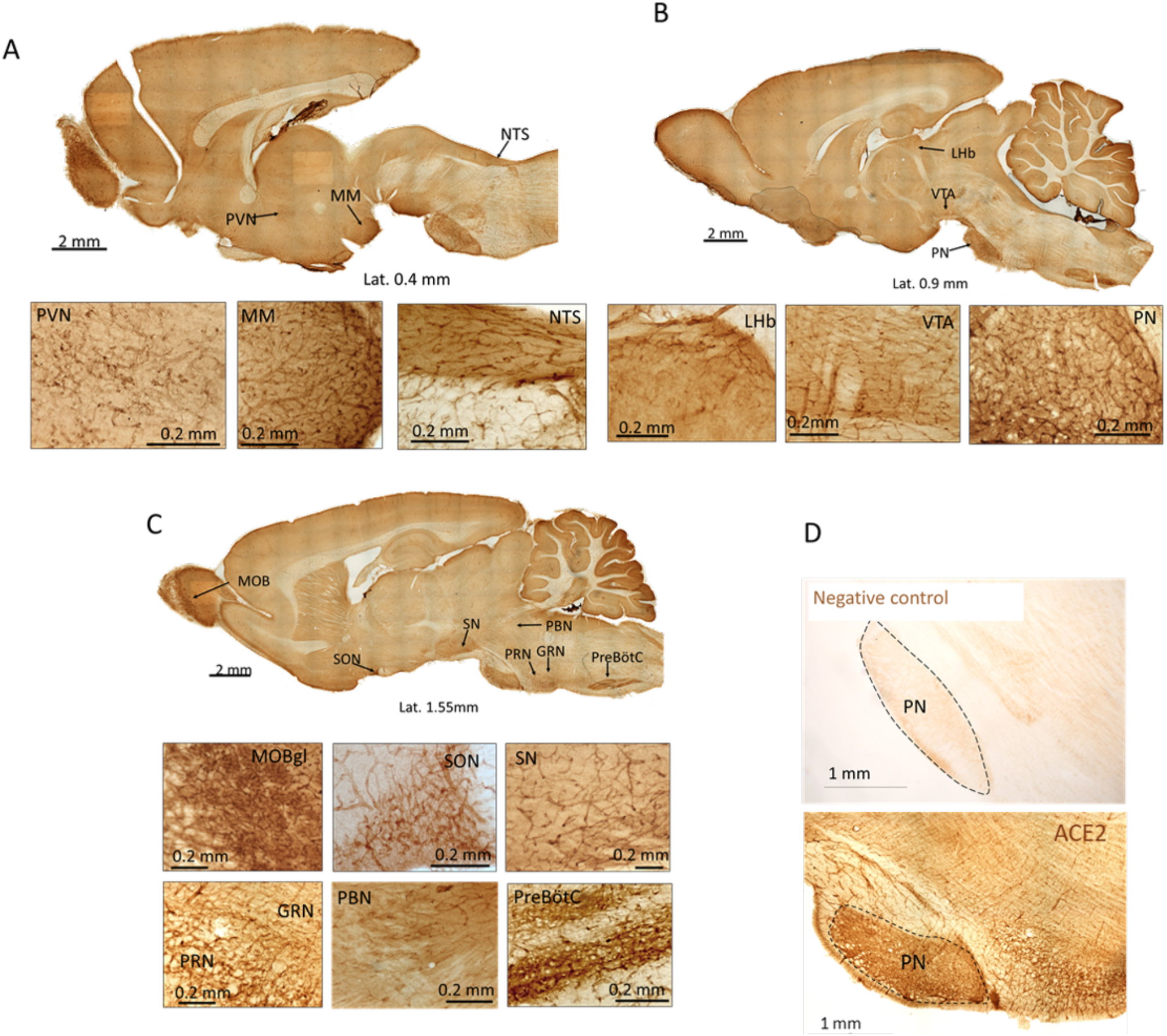
ACE2 expression in the whole rat brain. A-C: Scanned photomicrographs of representative sections immunoreacted against ACE2 at three medio-lateral coordinates. High expression of ACE2 was observed in the vasculature of the glomerular layer of the main olfactory bulb (MOBgl, panel C), paraventricular hypothalamic (PVN, panel A) and supraoptic (SON, panel C) nuclei, mammillary nucleus (MM, panel A), lateral habenula (LHb, panel B), ventral tegmental area (VTA, panel B), substantia nigra (SN, panel C), pontine nucleus (PN, panel B), pontine reticular nucleus (PRN, panels C), gigantoreticularis nucleus (GRN, panels C), parabrachial nucleus (PB, panel C), pre-Bötzinger complex (PreBötC, panel C), and nucleus of the tractus solitarius (NTS, panel A). D. top panel shows the negative control of the immunohistochemistry comparing with the positive control (low panel).

**Table 2.**
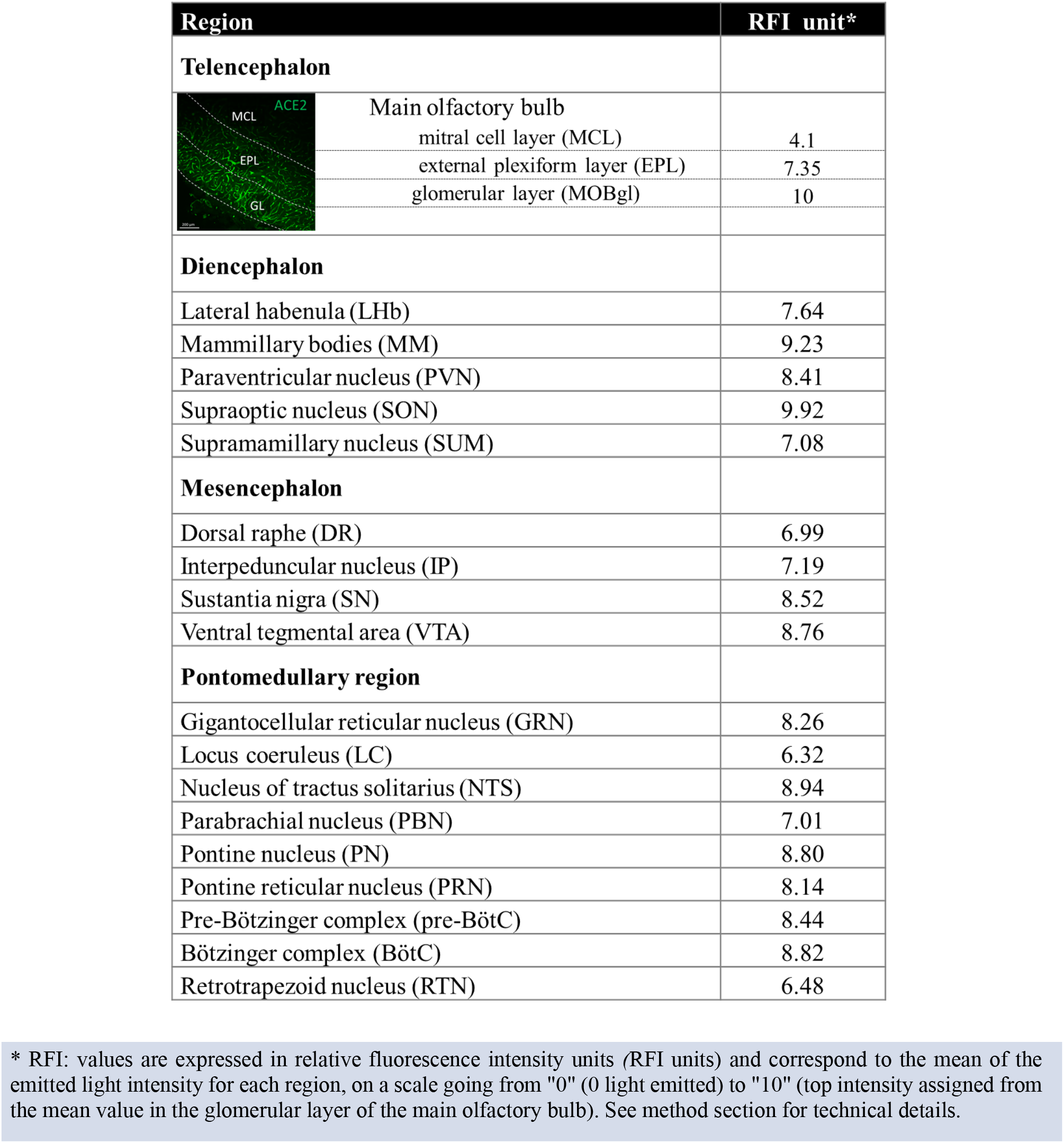
ACE2 immunoreactivity in richly vascularized regions of rat brain

### ACE2 expression in identified cells of the blood-brain barrier

Through GFAP immunoreaction and confocal microcopy assessment, we determined the expression of ACE2 in astrocytes throughout the brain. Some of those immunopositive cells were found extending their end-feet processes to surrounding blood vessels (Fig. 2A) or in close contact with the soma of neuronal bodies (Fig. 8). All identified pericytes also expressed ACE2, and some endothelial cells labelled with the antibody against Von Willebrand factor (VWF), also expressed ACE2 (Fig. 2B).

**Fig. 2.**
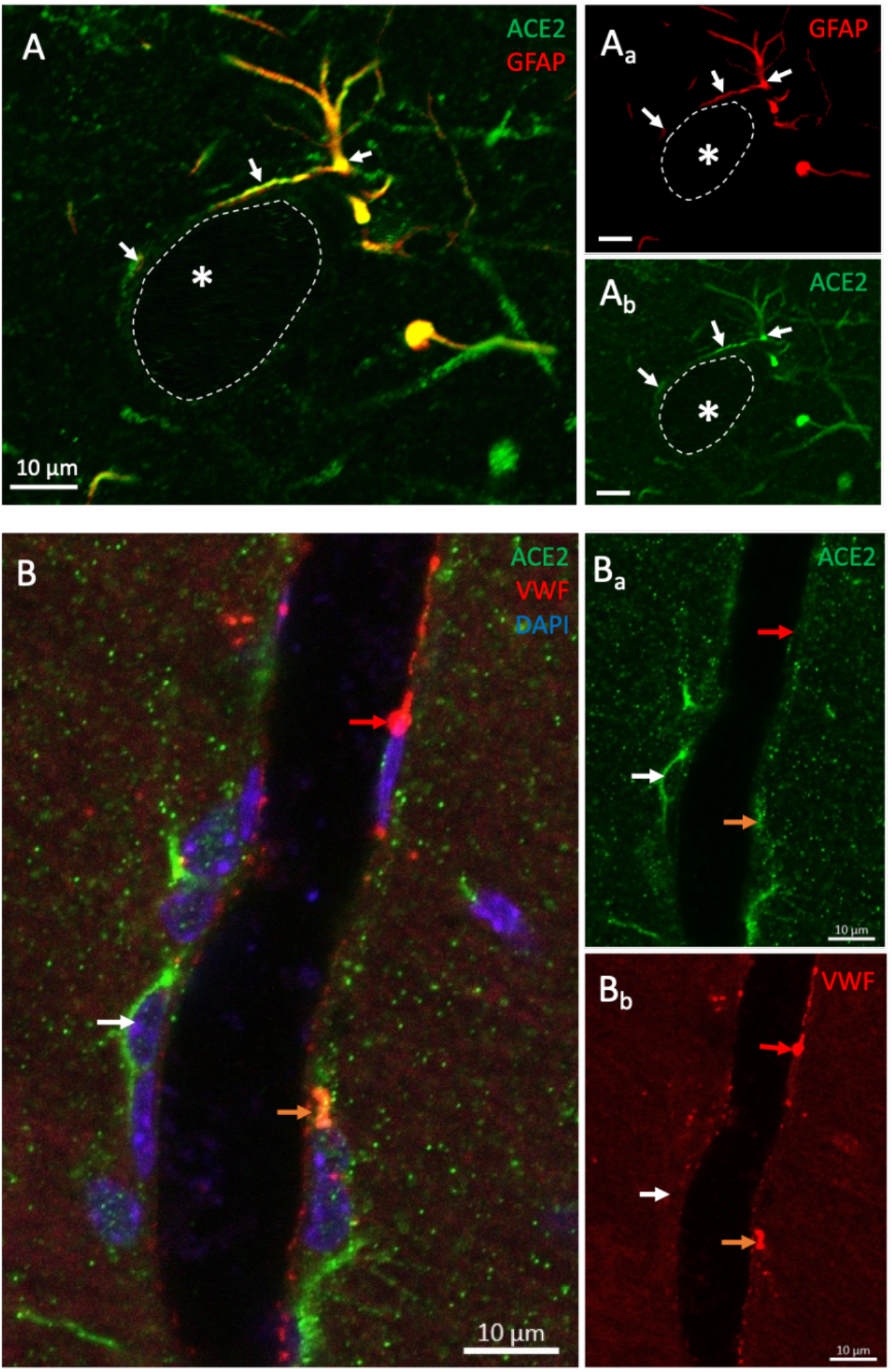
ACE2 is expressed in the components of the blood-brain barrier. A) Confocal micrographs show an astrocyte (GFAP labelling) positive for ACE2. Notice the ACE2 positive astrocytic processes surrounding a blood vessel delineated by dotted lines. B) Photomicrograph shows the expression of ACE2 in a pericyte (white arrow), and in an endothelial cell labelled for Von Willebrand factor (VWF) (orange arrow). An ACE2 negative endothelial cell is indicated by a red arrow.

**Fig. 8.**
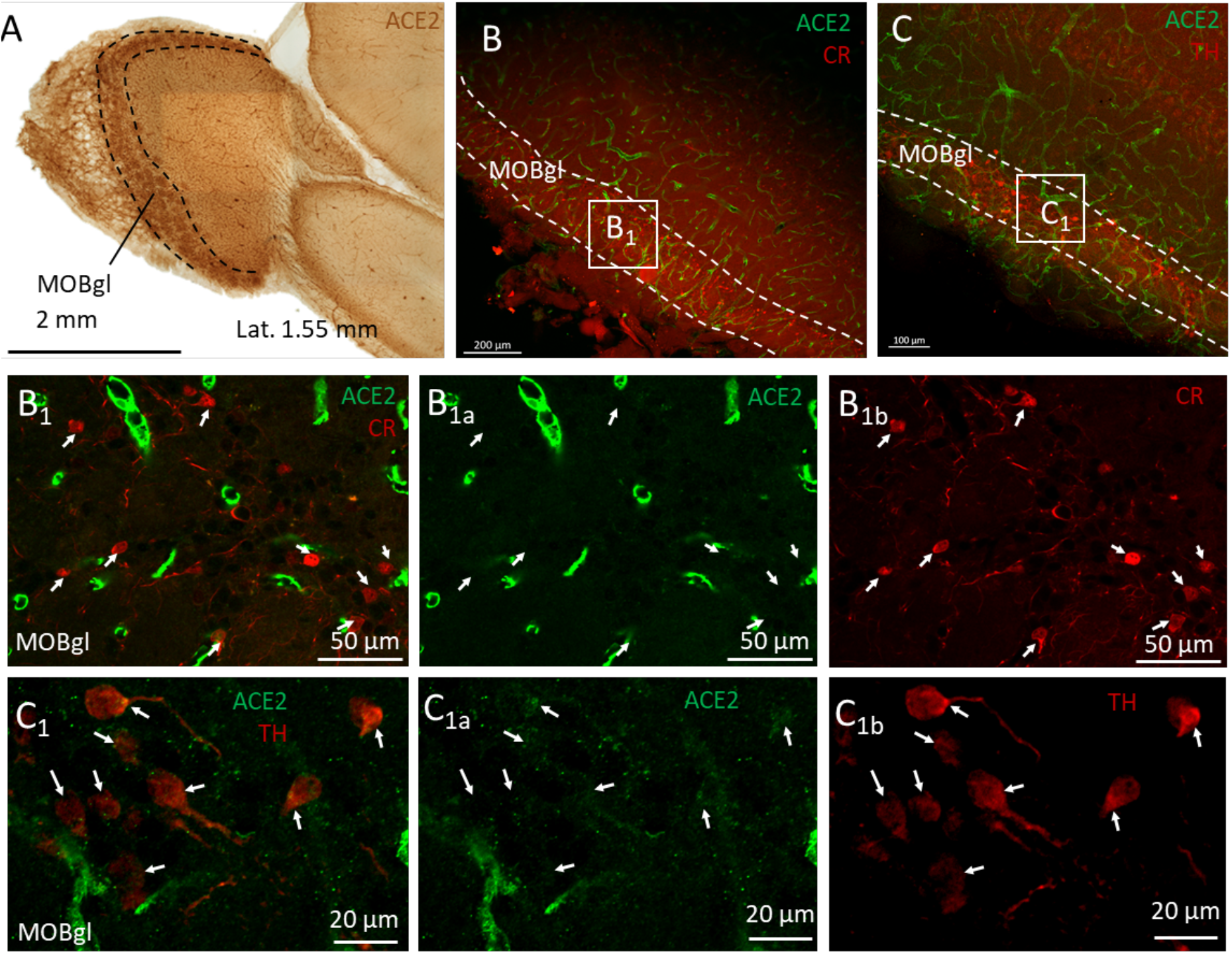
ACE2 is expressed in TH positive neurons of the main olfactory bulb (MOB). A) Low magnification photomicrograph of a DAB developed immunoreaction against ACE2 in the olfactory bulb region at the lateral 1.4 mm coordinate. The highest expression of ACE2 was found in the glomerular layer (MOBgl). Panels B and C show low power confocal photomicrographs of sections double labelled against ACE2 and calretinin (CR, panel B) or tyrosine hydroxylase (TH, panel C). High power photomicrographs of the squared regions labelled as B_1_ and C_1_ are shown in the lower panels. In the glomerular layer, ACE2 positive neurons were negative for CR (panel B_1s_) but positive for TH (panel C_1s_).

### ACE2 expression in brainstem arousal and respiratory networks

In the pontomedullary region, we found a high expression of ACE2 in the neuropil and in cells of the neurovascular unit clustered in nuclei that belong to the brainstem respiratory network (Fig. 3A). The nuclei where most of the ACE2 positive cells were observed were the parabrachial nucleus (PBN), nucleus of tractus solitarius (NTS), pre-Bötzinger complex (pre-BötC), retrotrapezoid nucleus (RTN), and Bötzinger complex. In the pre-BötC some of those neurons were further characterized as glycinergic, expressing the glycine transporter type 2 ((GlyT2 immunopositive), Fig. 3, inset of panel E).

**Fig.3.**
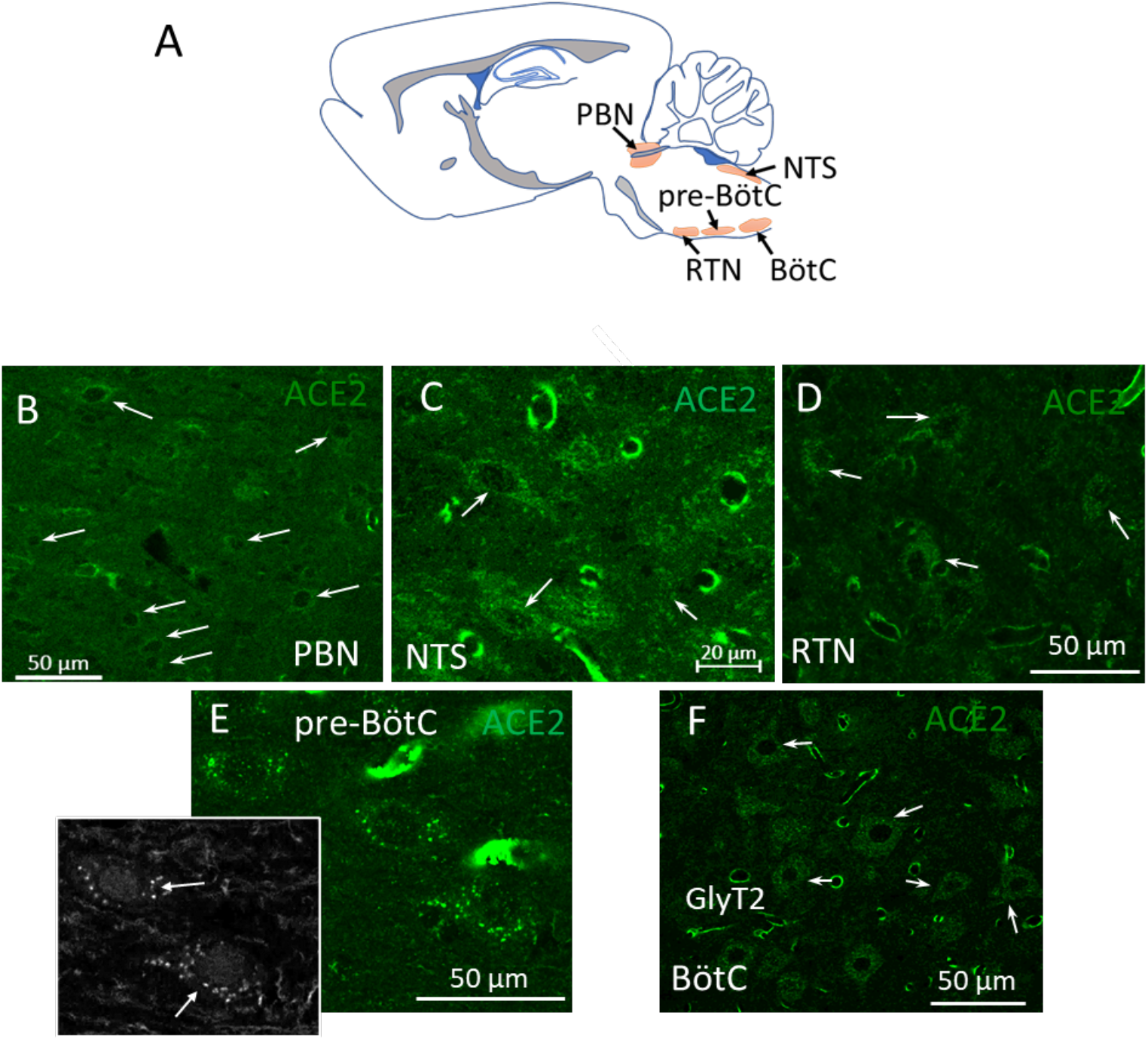
ACE2 is highly expressed in nuclei of the brainstem respiratory network. A: Schematic representation of the pontomedullary nuclei involved in breathing control, where high numbers of ACE2 expressing non-endothelial cells were found. B-F: High power confocal photomicrographs of the parabrachial nucleus (PBN), nucleus of tractus solitarius (NTS), retrotrapezoid nucleus (RTN), pre-Bötzinger complex (pre-BötC), and Bötzinger complex (BötC). Inset in panel E shows the colocalization of the glycine transporter 2 in glycinergic neurons of the pre-BötC. Arrows in all panels indicate some of the observed ACE2 positive cells.

In the arousal-related reticular formation, a high density of ACE2 positive cells was also found especially in the pontine reticular nucleus (PRN) and gigantocellular reticular nucleus (GRN), see figure 4.

**Fig. 4.**
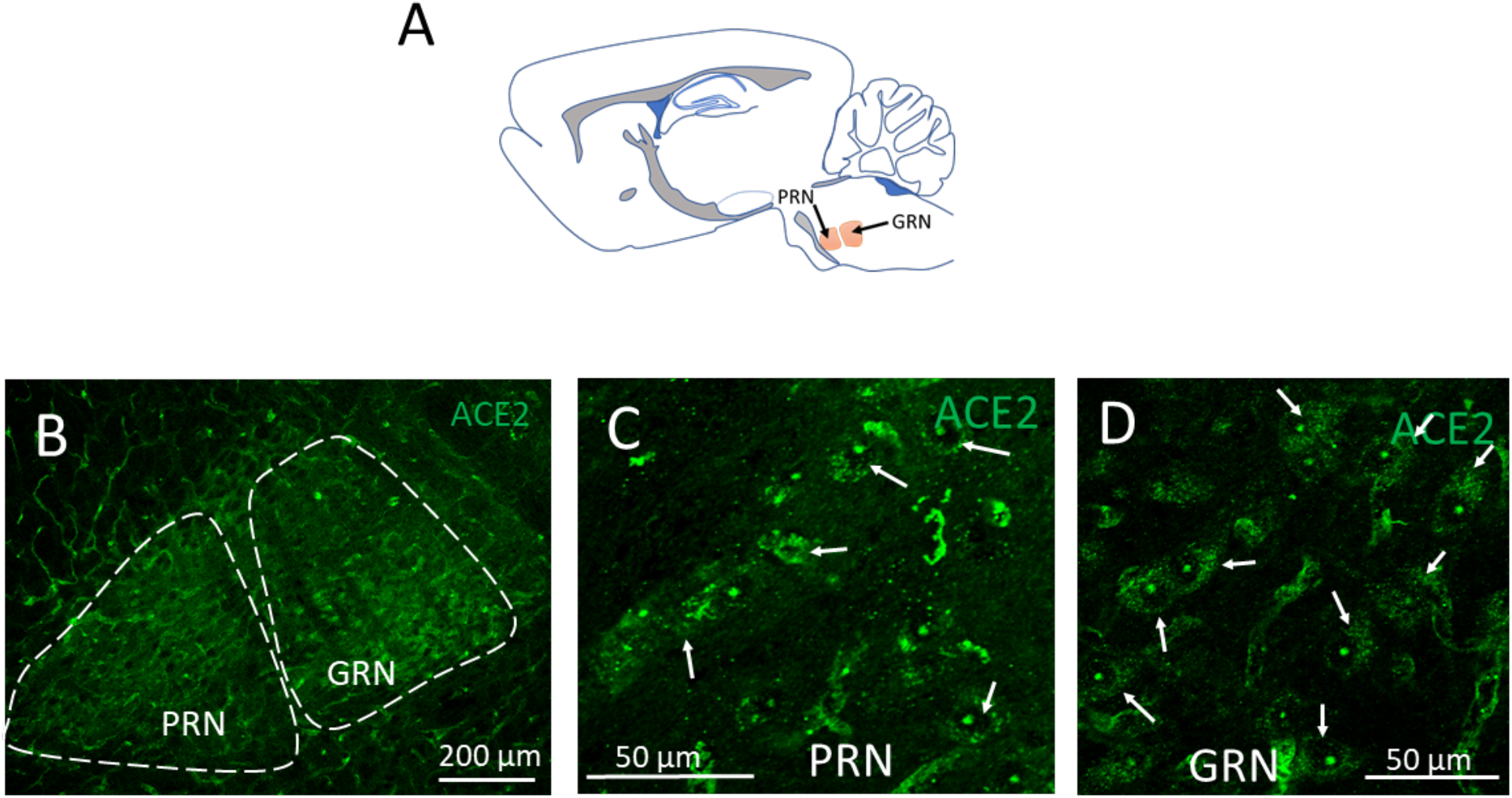
ACE2 is highly expressed in arousal-related reticular formation pontomedullary nuclei. A: Schematic representation of the location of the pontine reticular nuclei (PRN) and gigantocellular reticular nucleus (GRN) in a sagittal section. Low power (B) and high power (C and D) confocal photomicrographs show ACE2 positive cells identified in PRN. Arrows indicate examples of some of those cells.

### ACE2 expression in catecholaminergic and serotoninergic nuclei related to reward, movement and motivation networks

High ACE2 expression was observed in brainstem regions that contain dopaminergic, noradrenergic, and serotoninergic neurons. Figure 5 shows examples of ACE2-positive cells expressing tyrosine hydroxylase (TH), the rate-limiting enzyme for synthesis of dopamine and noradrenaline or tryptophan hydroxylase (TPH), the corresponding enzyme for the synthesis of serotonin. Double TH+ ACE2+ cells were found in the dopaminergic substantia nigra pars compacta (SNc, panels B) and ventral tegmental area (VTA, panels C) regions and in the noradrenergic region of locus coeruleus (LC). At the medio-lateral level 0.18 mm examined within the dorsal raphe (DR, panels E), we found a high density of TPH neurons co-expressing ACE2, the glycine transporter 2 (GlyT2), and calretinin (CR, a calcium binding protein), however we also found sparse TPH+ neurons that were negative for ACE2 and did not express GlyT2 or CR.

**Fig. 5.**
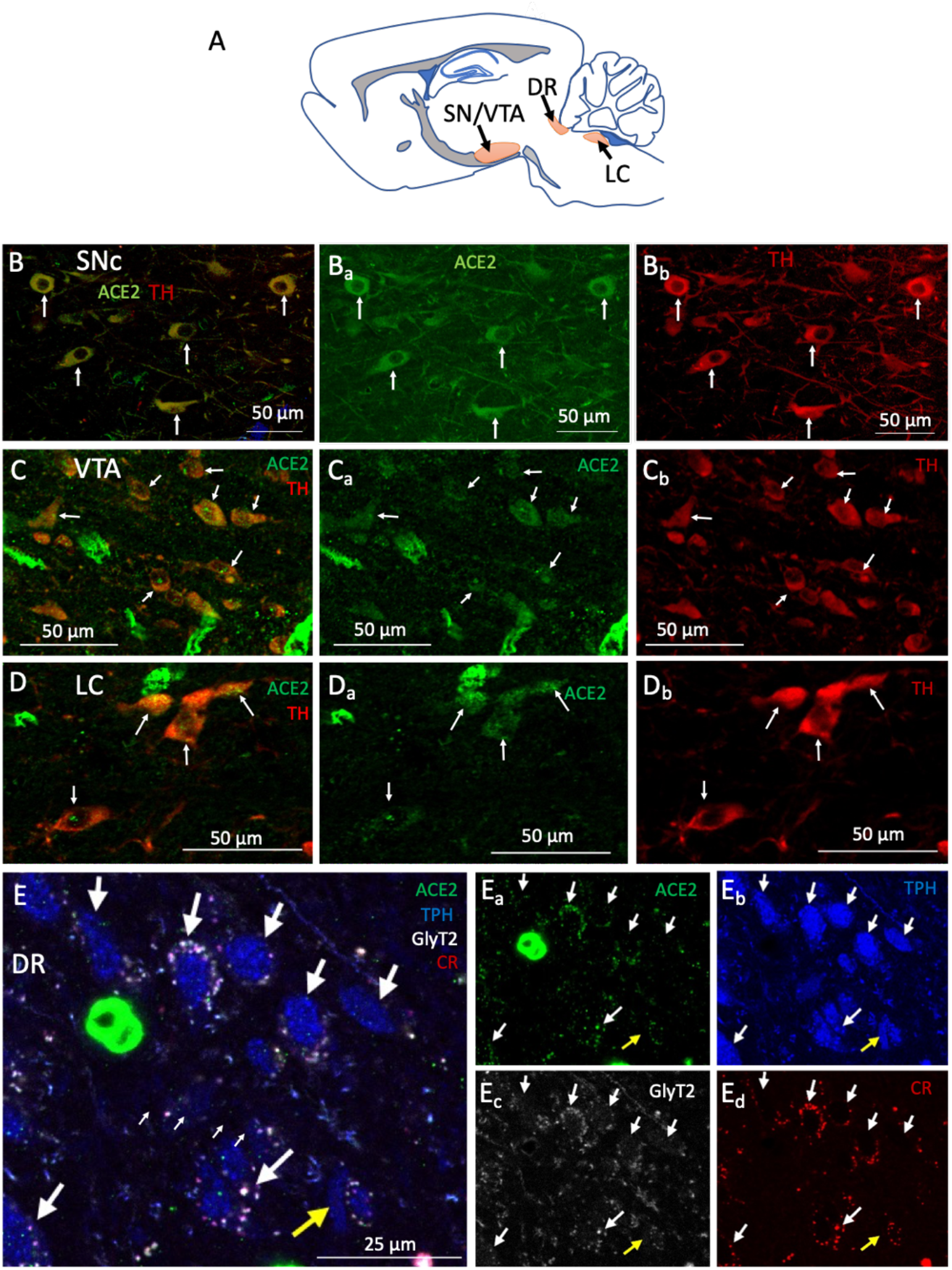
ACE2 is expressed in brainstem aminergic nuclei. A) Schematic representation of the evaluated brainstem aminergic nuclei location in a sagittal section. B - D) ACE2 is expressed in tyrosine hydroxylase (TH) expressing neurons within the substantia nigra pars compacta (SNc, panel B) and ventral tegmental area (VTA, panel C) dopaminergic regions, as well as in the locus coeruleus (LC, panel D) noradrenergic region. E) Many tryptophan hydroxylase (TPH), neurons in the serotoninergic dorsal raphe (DR) nucleus were positive for ACE2 and co-expressed the glycine transporter 2 (GlyT2) and the calcium binding protein calretinin (CR). Sparse TPH-positive neurons (indicated by the yellow arrow) do not express ACE2, CR nor GlyT2.

### ACE2 in diencephalic homeostatic networks

Compared to pontomedullary or mesencephalic ACE2 expression, lower ACE2 immunoactivity was observed in the diencephalon. However, we identified cells expressing ACE2 in the epithalamic lateral habenula (LHb) and in the hypothalamic paraventricular (PVN), supraoptic (SON) and suprammamillary (SUM) nuclei (figure 6). The lateral habenula, a key modulator of the aminergic centers of the midbrain, hosted a significant number of ACE2 positive for CR (figure 6, panels Bs). In PVN, we found that most of the vasopressin (AVP) expressing neurons co-express ACE2 (figure 6, panels Cs). In the SUM, ACE2 expression was observed in TH-positive neurons, but not in other NeuN-positive cells (figure 6, panels Ds)

**Fig. 6.**
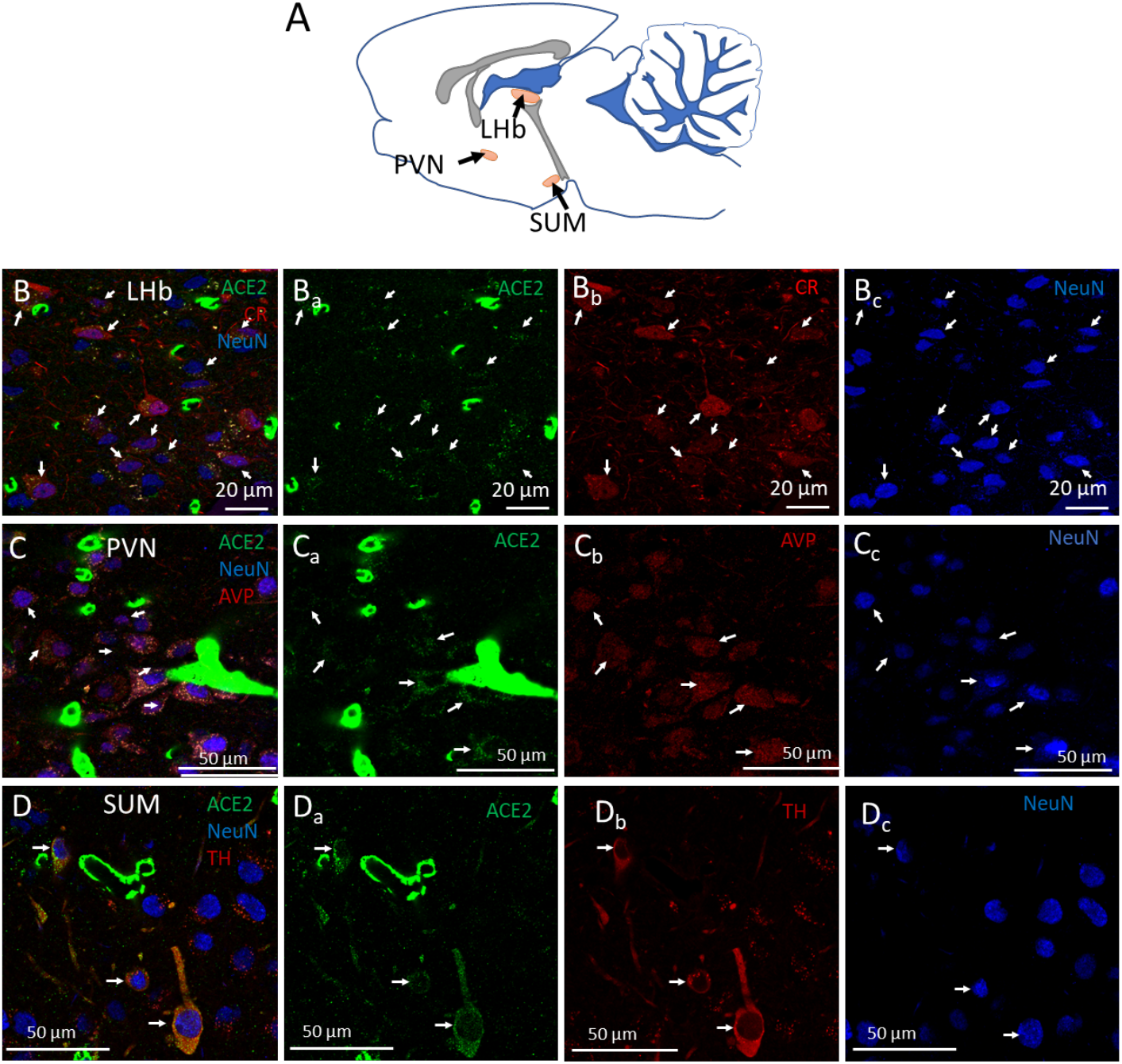
ACE2 is expressed in the diencephalic lateral habenula, paraventricular and suprammamillary neurons. A) Schematic representation in a sagittal slice, of the lateral habenula (LHb), paraventricular nucleus (PVN) and **supramammillary** nucleus (SUM), where some neurons expressing ACE2 were characterized. B) Confocal photomicrograph showing (arrows) that ACE2 is expressed in calretinin (CR) positive neurons (NeuN, Neuron-Specific Nuclear Protein). B) Arrows in photomicrographs show the colocalization of ACE2 in vasopressinergic (AVP) positive neurons (NeuN). D) In the SUM we found tyrosine hydroxylase (TH) positive neurons (NeuN) that co-express ACE2. Other non-TH neurons did not express ACE2.

### Expression of ACE2 in cognitive networks

The hippocampus has a high expression of ACE2 in astrocytes and in some neurons mainly located below the principal cell layers of CA1, and CA2. Astrocytic processes containing ACE2, appear to contact the soma of the principal neurons in all of these layers. Fig. 7 shows the three Cornu Ammonis (CA) areas with white arrows indicating identified interneurons expressing ACE2 and yellow arrows indicating the GFAP-expressing astrocytes in close proximity to the neuronal principal layer.

**Fig. 7.**
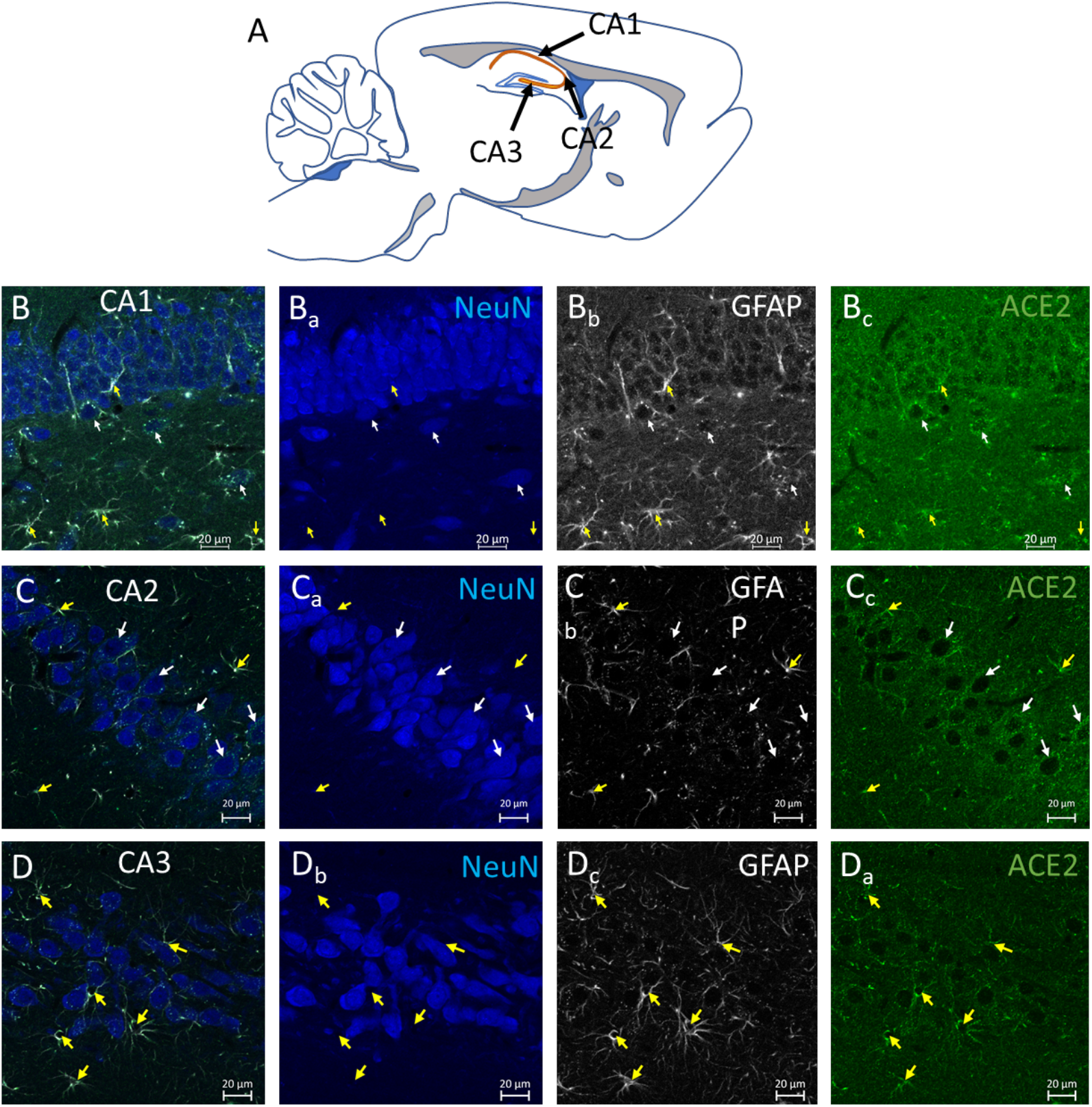
ACE2 is expressed in neurons and astrocytes in the hippocampus. A) Schematic representation of a sagittal brain slice, indicating the location of the CA (Cornu Ammonis) fields of hippocampus in orange color, where the high-power photomicrographs of the following panels were taken. B-D) Confocal photomicrographs in the regions of CA1 (panels Bs), CA2 (Panels Cs) and CA3 (Panels Ds), showing neurons that were double labelled with ACE2 and NeuN (white arrows) or astrocytes labelled with ACE2 and GFAP (yellow arrows). Note the presence of the GFAP+ /ACE+ astrocytic processes, projecting into the pyramidal layer. Also notice that the ACE2 positive neurons were mainly outside the pyramidal layer.

### Expression of ACE2 in the main olfactory bulb

The olfactory bulb is one of the regions with the highest expression of ACE2, especially in the glomerular layer (MOBgl) (Table 2). We investigated ACE2 co-expression with TH or CR, molecules known to be expressed in the glomerular layer. We found that ACE2 is expressed in TH-positive, but not CR-positive neurons in MOB (Fig. 8).

## Discussion

In this study, we report a widespread expression of ACE2 immunoreaction through the rat brain, in a cell type and anatomical regional specific manner. It is particularly remarkable that in the brain vasculature, the constituent elements for blood-brain barrier all expressed ACE2, with particularly high density of immunopositive vasculatures in the olfactory bulb, the hypothalamic nuclei, the midbrain dopaminergic regions and in various brainstem respiratory nuclei associated with breathing regulation and arousal. Co-expression of ACE2 with other molecular signatures in localized groups of neurons, participants of several well-defined circuits, such as for the respiratory rhythm, arousal, reward, homeostasis, learning, and memory are described here in detail. The potential impairment of these structures could be the neural substrates for the clinical manifestations of COVID-19 and the post-acute COVID-19 syndrome.

The expression of ACE2 in different tissues and organs has been associated with maintenance of diverse physiological processes, for instance, regulating cell survival, oxidative stress, angiogenesis, inflammation, and fibrosis (Hamming, Timens et al. 2004, Rabelo, Alenina et al. 2011, Uhal, Li et al. 2011). The first report on the brain expression of ACE2 was from tissues obtained from human autopsies (Harmer, Gilbert et al. 2002). Initially, ACE2 was thought to be present only in brain endothelial and smooth muscle cells (Hamming, Timens et al. 2004) albeit animal studies in rats showed that astrocytes also expressed ACE2 (Gallagher, Chappell et al. 2006). In another study, neuron-specific expression of ACE2 was reported (Doobay, Talman et al. 2007). Recent studies in human and mouse neuroblastoma, glioblastoma and microglial cell lines have also shown that ACE2 is expressed in all these cell lines, at both RNA and protein levels (Qiao, Li et al. 2020).

ACE2 was the putative receptor for severe acute respiratory syndrome (SARS) coronaviruses (Li, Moore et al. 2003) and SARS-CoV (Lan, Ge et al. 2020, Lu, Zhao et al. 2020). Moreover, coronaviruses can enter the brain and infect neurons and glial human cells (Arbour, Day et al. 2000, Cheng, Yang et al. 2020), and evidence of CNS coronavirus infection has been found in neural cells and cerebrospinal fluid (CSF) of COVID-19 patients (Arbour, Day et al. 2000, Cheng, Yang et al. 2020, Karvigh, Vahabizad et al. 2021, Song, Zhang et al. 2021). All of these observations, amidst the complexity of COVID-19, suggests that the regulation of ACE2 itself, could play a crucial role in COVID-19 pathophysiology including neurological symptoms and complications.

### ACE2 in the blood-brain barrier cellular components

The blood brain barrier (BBB) is crucial to maintain an adequate milieu for a normal physiological activity of neurons in different brain areas. It is composed of a continuous, non-fenestrated endothelium, and the accessory peri-endothelial structures as pericytes, astrocytes and a thin basement membrane (Reese and Karnovsky 1967). Endothelial cells through tight junctions seal the paracellular space, restricting the passage of substances to transport mediated by transporters or transcytosis. In the peri-endothelial space, astrocytes and pericytes regulate capillary flow and transcytosis (Armulik, Genove et al. 2010, Segarra, Aburto et al. 2021).

In this study, we showed that the density and distribution of immunopositive vasculature was heterogeneous between brain areas, with the highest density in the olfactory bulb, the supraoptic and paraventricular nuclei, and the mammillary bodies of the hypothalamus, the midbrain dopaminergic substantia nigra and ventral tegmental area, and nuclei in the pons and medulla involved in the regulation of breathing and arousal (Table2). Through immunofluorescence, we reported the expression of ACE2 in the cytoplasm of astrocytes, which are key components of the BBB. Notably, we noted that the endfoot processes of these cells were surrounding blood vessels (Fig. 2) and also had close contact with neurons (Fig. 7), suggesting a possible route for neuronal infection. The other cells of the BBB are the pericytes and the endothelial cells, we found that ACE2 was expressed in all pericytes and in some endothelial cells.

Recent histopathological studies from patients who died from severe COVID-19 indicate the presence of endothelial inflammation at gray matter structures (Kirschenbaum, Imbach et al. 2021)). Neuroinflammatory abnormalities during COVID-19 could be partially explained based on changes in ACE2 expression at the BBB of the vast brain network of capillaries, which could affect integrity of endothelial tight junctions allowing passage of cytokines and inflammatory cells into the brain. The diffusion of SARS-CoV-2 directly to nervous tissue and/or by infiltration of infected lymphocytes to the perivascular space, would leave glia and neurons directly exposed to the virus. Furthermore, cytokines might cause abnormalities in the excitability of neurons by affecting neurotransmitter release, cell survival and synaptic integrity in certain brain circuits, resulting in functional abnormalities (Vezzani and Viviani 2015).

It has been increasingly recognized that alterations in the normal brain hemodynamics is a common feature of COVID-19 and vascular events such as stroke are frequent (Kakarla, Kaneko et al. 2021, Sashindranath and Nandurkar 2021). Some imaging studies have shown alterations in cerebral blood flow in patients with COVID-19 (Sonkaya, OztUrk et al. 2020). And even in patients with no overt neurological manifestations, there are persistent alterations in the cerebral blood flow (Qin, Wu et al. 2021). The results we present in this study, on the vascular distribution of ACE2 in the brain, point out to potential regions where the presence of high levels of ACE2 in the vasculature, might serve as a docking point for SARS-CoV-2 entry into the brain.

### ACE2 in neuronal circuits

High-throughput single cell polymerase chain reaction and microarray studies have suggested the identity of cells or brain regions expressing ACE2 (Fodoulian, Tuberosa et al. 2020, Lukiw, Pogue et al. 2020, Muus, Luecken et al. 2021). However, little has been reported regarding the cellular identity of neurons expressing ACE2 or about the circuits where those neurons participate.

Here, we report that ACE2 is expressed in discrete neuronal groups throughout the brain, from the brainstem to the olfactory cortex. These neurons have a chemical signature that if altered, might cause functional abnormalities at different levels, and thus, explain some of the clinical manifestations of COVID-19. In the next sections we provide an analysis of the possible role of ACE2 expressing neurons in different circuits and the possible pathophysiological consequences of its disbalance in relevant physiological circuits.

### Circuits for respiratory control and arousal

Several nuclei involved in breathing and sleep-wake transitions are in the rhombencephalon, between the pons and medulla. The neurons responsible for modulating breathing can be largely found in three groups of nuclei located in the pons, the ventrolateral medulla, and the dorsolateral medulla. These nuclei are involved in the unconscious control of breathing by controlling the contraction and relaxation of respiratory muscles in response to signals from chemoreceptors that sense the concentration of oxygen or carbon monoxide in the blood (Dutschmann and Dick 2012). The pontine respiratory group is constituted by neurons in the parabrachial complex. Neurons in this group have been shown to project onto phrenic motoneurons and receive important sensory input from the nucleus of *tractus solitarius* (NTS) and chemoreceptor information from the RTN / PF nuclei. The activation of this nucleus causes abrupt cessation of inspiration and it has been implicated in resetting the respiratory rhythm in response to sensorial stimuli. Lesions to these nuclei inhibit the tachypnea induced by hypoxia or hypercapnia. The dorsal medullary group consist of neurons located in the NTS. At least two types of neurons have been identified, one type is presynaptic to phrenic motoneurons, the second type has been shown to receive input from stretch receptors in the lung, and send outputs to other groups in the respiratory network (Bautista, Pitts et al. 2014). The ventral medullary respiratory group consist of the RNT/PFZ nuclei, the pre-Bötzinger complex and the Bötzinger complex. The RTN/PFZ has been shown to contain PHOX2B-positive neurons able to sense CO2 concentrations (Onimaru, Ikeda et al. 2014). This area also receives information about oxygen levels from the carotid bodies (Guyenet, Stornetta et al. 2019). Mutations in PHOX2B are associated with a syndrome of congenital hypoventilation in humans (Amiel, Laudier et al. 2003). The Bötzinger complex has mainly an expiratory function with the adjacent pre-Bötzinger complex involved mainly in the inspiratory phase of breathing (Smith, Abdala et al. 2009). The Bötzinger complex contains a large population of glycinergic neurons (Winter, Fresemann et al. 2009). The presence of angiotensin II in the respiratory network was noted in the late eighties by Aguirre et al, who reported a cluster of angiotensin II immunopositive somata in the Bötzinger complex and Parabrachial nucleus and abundant immunopositive fibers in the nucleus tractus solitarius (NTS) and medial PBN (Aguirre, Covenas et al. 1989). However, the exact chemosensory mechanism of this process remained elusive. A recent study used a microarray design to quantify the expression of ACE2 RNA in some samples of human brain autopsies, the results showed that the most intense expression of ACE2 was in the medulla and pons. In this study, we report the presence of a large group of neurons, located in different dorsal and ventral nuclei of the respiratory complex that are ACE2 immunopositive (figure 3). From the pontine respiratory group, we found ACE2 expression in neurons of the parabrachial nucleus (PBN). From the ventral respiratory group, ACE2 was expressed in neurons of the retrotrapezoid nucleus (RTN), in the glycinergic transporter (GlyT2) expressing neurons of the pre-Bötzinger complex (PreBötC) and in the Bötzinger complex (BötC). From the dorsal respiratory group, we found ACE2 positive cells in NTS. From the reticular formation, ACE2 positive cells were identified in the caudal pontine reticular nucleus (PRN) and gigantocellular reticular nucleus (GRN) (Fig. 3).

One of the main symptoms of COVID-19 patients is the lack of perception of dangerously low levels of blood oxygen, a phenomenon called “happy hypoxemia”(U and Verma 2020). Also of great concern is the high incidence of patients unable to restore their normal autonomic ventilation after having been under artificially controlled ventilation. A possible hypothesis to explain these observations, is that SARS-CoV-2 exerts cytotoxic effects on the infected cells of the respiratory network. This could impede restoration of normal ventilatory rhythmicity or affect chemosensation of CO2 and O2 levels in blood, thus inhibiting the perception of dangerously levels of hypoxia or hypercapnia. A final possibility to explain the ventilatory anomalies observed in COVID-19 patients, is that infection of ACE2-expressing cells down-regulates expression of the latter, reducing the transformation of angiotensin II into angiotensin I. Excessive angiotensin II could affect the function of circuits involved in breathing control. In support of this hypothesis, an association between genetic variation in ACE activity, and fatal cardiorespiratory pathology, has been reported (Harding, Dhamrait et al. 2003).

### Circuits for hydroelectrolytic homeostasis

The paraventricular (PVN) and supraoptic (SON) nuclei of the hypothalamus are crucial to regulate blood volume and osmolarity (Dunn, Brennan et al. 1973). However, these nuclei also innervate central targets in limbic and brainstem regions such as the hippocampus, amygdala, habenula, and locus coeruleus and modulate behavioral cognitive and emotional responses (Cui, Gerfen et al. 2013, Zhang and Hernandez 2013, Hernandez, Vazquez-Juarez et al. 2015, Hernandez, Hernández et al. 2016, Hernandez, Hernandez et al. 2016, Zhang, Hernandez et al. 2016).

We found that PVN and SON are richly vascularized structures in the brain with high ACE2 expression. Any vascular disruption would necessarily affect the normal functioning of those nuclei. Moreover, we found ACE2 expression also in vasopressinergic neurons, confirming previous reports using transcriptomic analysis (Nampoothiri, Sauve et al. 2020). It is known that the secretion of vasopressin is regulated by the RAAS. Angiotensin II stimulates the release of vasopressin (Sandgren, Linggonegoro et al. 2018). Mice overexpressing ACE2 showed a diminished hypertensive response to the systemic administration of angiotensin II (Feng, Hans et al. 2012) and reduced deoxycorticosterone acetate (DOCA)-salt induced hypertension (Xia, de Queiroz et al. 2015). Also, pharmacological interventions that increase ACE2 expression in the brain, have been shown to produce antihypertensive effects and a decrease in vasopressin release (Hmazzou, Marc et al. 2021). A series of case reports have noted the presence of syndrome of inadequate antidiuretic hormone secretion (SIADH) in COVID-19 related pneumonia (Yousaf, Al-Shokri et al. 2020). Current literature suggests that the increase in vasopressin seen in some patients with COVID-19 is mediated by systemic inflammation, which acts as a non-osmotic stimuli for vasopressin production (Swart, Hoorn et al. 2011) and kidney damage (Gheorghe, Ilie et al. 2021). However, it could be argued that SARS-CoV2 directly invades the vascular structures that nourish important AVP secreting nuclei in the brain, such as the PVN and SON, thus causing important disruption in homeostasis that leads to altered hormonal secretion and, consequently, SIADH, or that the direct infection of AVP producing neurons by SARS-CoV-2, leads to an internalization of its receptor and subsequent accumulation of the hormone angiotensin II stimulatory for AVP-release.

### Circuits for reward, motivation, and movement

The two main dopaminergic populations of the midbrain are the ventral tegmental area (VTA) and the *sustantia nigra pars compacta* (SNpc). Three main pathways arise from these structures. The ascending projections from the VTA to the nucleus accumbens are called the mesolimbic pathway and have classically been described as part of a reward system (Wise 1978, Schultz 2002). The projections from VTA to the prefrontal cortex are called the mesocortical pathway and have been associated with modulation of cognitive, working memory and decision-making functions (Tanaka 2006, Lapish, Kroener et al. 2007, Verharen, de Jong et al. 2018). Finally, the nigrostriatal pathway arises in the SN and terminates in the caudate and putamen, it has been associated with the modulation of voluntary movement (Smith and Bolam 1990, Prensa, Gimenez-Amaya et al. 2009). These circuits are modulated by excitatory projections from the lateral habenula (LHb), via an inhibitory relay in the tail of the VTA (Barrot, Sesack et al. 2012, Bourdy and Barrot 2012, Stamatakis and Stuber 2012). In this study, we found ACE2/tyrosine hydroxylase (TH) co-expressing neurons in both VTA and SNpc.

The dorsal raphe nucleus (DRN) hosts the largest group of serotoninergic neurons and have classically been implicated in modulation of mood. Studies have shown alterations in anatomy or function of this nucleus in depression or anxiety and in suicide patients (Underwood, Khaibulina et al. 1999, Bruschetta, Jin et al. 2020, Li, Zhou et al. 2020). This nucleus also participate in modulating the core respiratory networks of the brainstem (Bautista, Pitts et al. 2014). The lateral habenula have reciprocal connections with the DRN and is a critical modulator of the activity of serotoninergic neurons. This LHb-DRN circuit has been implicated in the regulation of cognition, reward, and arousal functions (Zhao, Zhang et al. 2015). In this study, we found ACE2/tryptophan hydroxylase (TPH) co-expressing neurons in the DRN, some of them co-express the GlyT2, which is an important population of projection neurons from DRN (Krzywkowski, Jacobowitz et al. 1995).

Interactions between brain*-RAAS* (McKinley, Albiston et al. 2003, Huang and Leenen 2009, Cosarderelioglu, Nidadavolu et al. 2020) and the dopaminergic system have been reported. The infusion of angiotensin II in striatum provoke local dopamine release (Brown, Steward et al. 1996). Angiotensin II has been shown to regulate the synthesis of the enzymes involved in catecholamine biosynthesis (Aschrafi, Berndt et al. 2019). Moreover, ACE2 has been reported in the mitochondria isolated form cell cultures derived from dopaminergic neurons (Costa-Besada, Valenzuela et al. 2018). It is normally considered that SARS-CoV-2 induce a downregulation of its receptor (Kuba, Imai et al. 2005, Peiro and Moncada 2020, Verdecchia, Cavallini et al. 2020, Zhang, Zetter et al. 2021). Diminishing the activity of ACE2 would lead to accumulation of angiotensin II and possibly leading to a dysregulation of dopaminergic and serotoninergic neurons expressing ACE2. This might partly explain emotional, cognitive, motivational and locomotor symptoms observed in COVID-19. Moreover, the increased incidence of Parkinson and parkinsonism reported in the recent literature (Helmich and Bloem 2020, Leta, Rodriguez-Violante et al. 2021, Morassi, Palmerini et al. 2021) could be related to the above catecholamine metabolic disfunction.

### Circuits for sensory processing

In this work we found ACE2 expression in the glomerular layer of the main olfactory bulb (MOBgl), the enzyme was present in TH positive neurons but not in calretinin positive neurons (Fig. 8). The olfactory bulb contains the largest population of dopaminergic neurons in the rat brain, and those have been identified mainly as periglomerular, and playing a role in the codification of the olfactory processing (Pignatelli and Belluzzi 2017, Kosaka, Pignatelli et al. 2020). Also, ACE2 neurons were observed in the nucleus of the tractus solitarius (NTS) and in the upstream nuclei in the gustatory pathway, the parabrachial nucleus (PBN). Besides their role in the processing of taste information, these nuclei, as discussed above also participate in the processing of chemosensory information related to the sensing of oxygen and CO2 levels. One of the most classical symptoms of COVID-19 is the loss of taste and smell (Butowt and von Bartheld 2020, Mao, Jin et al. 2020). It may be possible that dysfunction of ACE2 expressing neurons in these sensorial circuits underly the anosmia and ageusia observed in the acute phase of the disease, and that long lasting changes in these circuits might explain the protracted alterations that some patients report after recovery (Marshall 2021).

The distribution of ACE2 in neuronal, astrocytic, and epithelial cells of the mammalian central nervous system, especially the evidence for heterogeneity in abundance of expression throughout the brain, provides a template for understanding neurological manifestations of SARS-CoV-2 infection both during infection and following viral clearance. Changes in the abundance of ACE2, its effect on the peptides generated by this enzyme, as well as selective neuronal damage/death in a given neural circuit, merit urgent further investigation.

## Acknowledgements

We thank Dr. Rafael Hernández for providing animal facility assistance during the public health emergency. MAZ thanks postdoctoral fellowship through the ALIANZA UCMX / Innova UNAM grant #013-2020. OH-P is a postdoctoral fellowship from Programa de Becas Posdoctorales of DGAPA/UNAM. This study was supported by grants: UNAM-DGAPA-PAPIIT-GI200121 & CONACYT-CB-238744 (LZ); NIMH-IRP ZIAMH002498 (LEE).

## REFERENCES

Aguirre, J. A., R. Covenas, D. Croix, J. R. Alonso, J. A. Narvaez, G. Tramu and S. Gonzalez-Baron (1989). “Immunocytochemical study of angiotensin-II fibres and cell bodies in the brainstem respiratory areas of the cat.” Brain Res 489(2): 311–317.

Amiel, J., B. Laudier, T. Attie-Bitach, H. Trang, L. de Pontual, B. Gener, D. Trochet, H. Etchevers, P. Ray, M. Simonneau, M. Vekemans, A. Munnich, C. Gaultier and S. Lyonnet (2003). “Polyalanine expansion and frameshift mutations of the paired-like homeobox gene PHOX2B in congenital central hypoventilation syndrome.” Nat Genet 33(4): 459–461.

Arbour, N., R. Day, J. Newcombe and P. J. Talbot (2000). “Neuroinvasion by human respiratory coronaviruses.” J Virol 74(19): 8913–8921.

Armulik, A., G. Genove, M. Mae, M. H. Nisancioglu, E. Wallgard, C. Niaudet, L. He, J. Norlin, P. Lindblom, K. Strittmatter, B. R. Johansson and C. Betsholtz (2010). “Pericytes regulate the blood-brain barrier.” Nature 468(7323): 557–561.

Aschrafi, A., A. Berndt, J. A. Kowalak, J. R. Gale, A. E. Gioio and B. B. Kaplan (2019). “Angiotensin II mediates the axonal trafficking of tyrosine hydroxylase and dopamine beta-hydroxylase mRNAs and enhances norepinephrine synthesis in primary sympathetic neurons.” J Neurochem 150(6): 666–677.

Barrot, M., S. R. Sesack, F. Georges, M. Pistis, S. Hong and T. C. Jhou (2012). “Braking dopamine systems: a new GABA master structure for mesolimbic and nigrostriatal functions.” J Neurosci 32(41): 14094–14101.

Bautista, T. G., T. E. Pitts, P. M. Pilowsky and K. F. Morris (2014). Chapter 18 - The Brainstem Respiratory Network. Neuronal Networks in Brain Function, CNS Disorders, and Therapeutics. C. L. Faingold and H. Blumenfeld. San Diego, Academic Press: 235–245.

Bourdy, R. and M. Barrot (2012). “A new control center for dopaminergic systems: pulling the VTA by the tail.” Trends Neurosci 35(11): 681–690.

Brown, D. C., L. J. Steward, J. Ge and N. M. Barnes (1996). “Ability of angiotensin II to modulate striatal dopamine release via the AT1 receptor in vitro and in vivo.” Br J Pharmacol 118(2): 414–420.

Bruschetta, G., S. Jin, Z. W. Liu, J. D. Kim and S. Diano (2020). “MC4R Signaling in Dorsal Raphe Nucleus Controls Feeding, Anxiety, and Depression.” Cell Rep 33(2): 108267.

Butowt, R. and C. S. von Bartheld (2020). “Anosmia in COVID-19: Underlying Mechanisms and Assessment of an Olfactory Route to Brain Infection.” Neuroscientist: 1073858420956905.

Cheng, Q., Y. Yang and J. Gao (2020). “Infectivity of human coronavirus in the brain.” EBioMedicine 56: 102799.

Cohen, M. E., R. Eichel, B. Steiner-Birmanns, A. Janah, M. Ioshpa, R. Bar-Shalom, J. J. Paul, H. Gaber, V. Skrahina, N. M. Bornstein and G. Yahalom (2020). “A case of probable Parkinson’s disease after SARS-CoV-2 infection.” Lancet Neurol 19(10): 804–805.

Cosarderelioglu, C., L. S. Nidadavolu, C. J. George, E. S. Oh, D. A. Bennett, J. D. Walston and P. M. Abadir (2020). “Brain Renin-Angiotensin System at the Intersect of Physical and Cognitive Frailty.” Front Neurosci 14: 586314.

Costa-Besada, M. A., R. Valenzuela, P. Garrido-Gil, B. Villar-Cheda, J. A. Parga, J. L. Lanciego and J. L. Labandeira-Garcia (2018). “Paracrine and Intracrine Angiotensin 1-7/Mas Receptor Axis in the Substantia Nigra of Rodents, Monkeys, and Humans.” Mol Neurobiol 55(7): 5847–5867.

Cui, Z., C. R. Gerfen and W. S. Young, 3rd (2013). “Hypothalamic and other connections with dorsal CA2 area of the mouse hippocampus.” J Comp Neurol 521(8): 1844–1866.

Ding, Q., N. V. Shults, B. T. Harris and Y. J. Suzuki (2020). “Angiotensin-converting enzyme 2 (ACE2) is upregulated in Alzheimer’s disease brain.” bioRxiv.

Donoghue, M., F. Hsieh, E. Baronas, K. Godbout, M. Gosselin, N. Stagliano, M. Donovan, B. Woolf, K. Robison, R. Jeyaseelan, R. E. Breitbart and S. Acton (2000). “A novel angiotensin-converting enzyme-related carboxypeptidase (ACE2) converts angiotensin I to angiotensin 1-9.” Circ Res 87(5): E1–9.

Doobay, M. F., L. S. Talman, T. D. Obr, X. Tian, R. L. Davisson and E. Lazartigues (2007). “Differential expression of neuronal ACE2 in transgenic mice with overexpression of the brain renin-angiotensin system.” Am J Physiol Regul Integr Comp Physiol 292(1): R373–381.

Dunn, F. L., T. J. Brennan, A. E. Nelson and G. L. Robertson (1973). “The role of blood osmolality and volume in regulating vasopressin secretion in the rat.” J Clin Invest 52(12): 3212–3219.

Dutschmann, M. and T. E. Dick (2012). “Pontine mechanisms of respiratory control.” Compr Physiol 2(4): 2443–2469.

Feng, Y., C. Hans, E. McIlwain, K. J. Varner and E. Lazartigues (2012). “Angiotensin-converting enzyme 2 over-expression in the central nervous system reduces angiotensin-II-mediated cardiac hypertrophy.” PLoS One 7(11): e48910.

Fodoulian, L., J. Tuberosa, D. Rossier, M. Boillat, C. Kan, V. Pauli, K. Egervari, J. A. Lobrinus, B. N. Landis, A. Carleton and I. Rodriguez (2020). “SARS-CoV-2 Receptors and Entry Genes Are Expressed in the Human Olfactory Neuroepithelium and Brain.” iScience 23(12): 101839.

Gallagher, P. E., M. C. Chappell, C. M. Ferrario and E. A. Tallant (2006). “Distinct roles for ANG II and ANG-(1-7) in the regulation of angiotensin-converting enzyme 2 in rat astrocytes.” Am J Physiol Cell Physiol 290(2): C420–426.

Gheorghe, G., M. Ilie, S. Bungau, A. M. P. Stoian, N. Bacalbasa and C. C. Diaconu (2021). “Is There a Relationship between COVID-19 and Hyponatremia?” Medicina (Kaunas) 57(1).

Guyenet, P. G., R. L. Stornetta, G. Souza, S. B. G. Abbott, Y. Shi and D. A. Bayliss (2019). “The Retrotrapezoid Nucleus: Central Chemoreceptor and Regulator of Breathing Automaticity.” Trends Neurosci 42(11): 807–824.

Haidar, M. A., H. Jourdi, Z. Haj Hassan, O. Ashekyan, M. Fardoun, Z. Wehbe, D. Maaliki, M. Wehbe, S. Mondello, S. Abdelhady, S. Shahjouei, M. Bizri, Y. Mechref, M. S. Gold, G. Dbaibo, H. Zaraket, A. H. Eid and F. Kobeissy (2021). “Neurological and Neuropsychological Changes Associated with SARS-CoV-2 Infection: New Observations, New Mechanisms.” Neuroscientist: 1073858420984106.

Hamming, I., W. Timens, M. L. Bulthuis, A. T. Lely, G. Navis and H. van Goor (2004). “Tissue distribution of ACE2 protein, the functional receptor for SARS coronavirus. A first step in understanding SARS pathogenesis.” J Pathol 203(2): 631–637.

Harding, D., S. Dhamrait, N. Marlow, A. Whitelaw, S. Gupta, S. Humphries and H. Montgomery (2003). “Angiotensin-converting enzyme DD genotype is associated with worse perinatal cardiorespiratory adaptation in preterm infants.” J Pediatr 143(6): 746–749.

Harmer, D., M. Gilbert, R. Borman and K. L. Clark (2002). “Quantitative mRNA expression profiling of ACE 2, a novel homologue of angiotensin converting enzyme.” FEBS Lett 532(1-2): 107–110.

Helmich, R. C. and B. R. Bloem (2020). “The Impact of the COVID-19 Pandemic on Parkinson’s Disease: Hidden Sorrows and Emerging Opportunities.” J Parkinsons Dis 10(2): 351–354.

Hernandez, V., O. Hernández, M. Gomora, M. Perez De La Mora, K. Fuxe, L. Eiden and L. Zhang (2016). “Hypothalamic vasopressinergic projections innervate central amygdala GABAergic neurons: implications for anxiety and stress coping.” Frontiers in Neural Circuits 10(92).

Hernandez, V. S., O. R. Hernandez, M. Perez de la Mora, M. J. Gomora, K. Fuxe, L. E. Eiden and L. Zhang (2016). “Hypothalamic Vasopressinergic Projections Innervate Central Amygdala GABAergic Neurons: Implications for Anxiety and Stress Coping.” Front Neural Circuits 10: 92.

Hernandez, V. S., E. Vazquez-Juarez, M. M. Marquez, F. Jauregui-Huerta, R. A. Barrio and L. Zhang (2015). “Extra-neurohypophyseal axonal projections from individual vasopressin-containing magnocellular neurons in rat hypothalamus.” Front Neuroanat 9: 130.

Hmazzou, R., Y. Marc, A. Flahault, R. Gerbier, N. De Mota and C. Llorens-Cortes (2021). “Brain ACE2 activation following brain aminopeptidase A blockade by firibastat in salt-dependent hypertension.” Clin Sci (Lond) 135(6): 775–791.

Huang, B. S. and F. H. Leenen (2009). “The brain renin-angiotensin-aldosterone system: a major mechanism for sympathetic hyperactivity and left ventricular remodeling and dysfunction after myocardial infarction.” Curr Heart Fail Rep 6(2): 81–88.

Kabbani, N. and J. L. Olds (2020). “Does COVID19 Infect the Brain? If So, Smokers Might Be at a Higher Risk.” Mol Pharmacol 97(5): 351–353.

Kakarla, V., N. Kaneko, M. Nour, K. Khatibi, F. Elahi, D. S. Liebeskind and J. D. Hinman (2021). “Pathophysiologic mechanisms of cerebral endotheliopathy and stroke due to Sars-CoV-2.” J Cereb Blood Flow Metab: 271678X20985666.

Karvigh, S. A., F. Vahabizad, M. S. Mirhadi, G. Banihashemi and M. Montazeri (2021). “COVID-19-related refractory status epilepticus with the presence of SARS-CoV-2 (RNA) in the CSF: a case report.” Neurol Sci.

Kirschenbaum, D., L. L. Imbach, E. J. Rushing, K. B. M. Frauenknecht, D. Gascho, B. V. Ineichen, E. Keller, S. Kohler, M. Lichtblau, R. R. Reimann, K. Schreib, S. Ulrich, P. Steiger, A. Aguzzi and K. Frontzek (2021). “Intracerebral endotheliitis and microbleeds are neuropathological features of COVID-19.” Neuropathol Appl Neurobiol 47(3): 454–459.

Kosaka, T., A. Pignatelli and K. Kosaka (2020). “Heterogeneity of tyrosine hydroxylase expressing neurons in the main olfactory bulb of the mouse.” Neurosci Res 157: 15–33.

Krzywkowski, P., D. M. Jacobowitz and Y. Lamour (1995). “Calretinin-containing pathways in the rat forebrain.” Brain Res 705(1-2): 273–294.

Kuba, K., Y. Imai, S. Rao, H. Gao, F. Guo, B. Guan, Y. Huan, P. Yang, Y. Zhang, W. Deng, L. Bao, B. Zhang, G. Liu, Z. Wang, M. Chappell, Y. Liu, D. Zheng, A. Leibbrandt, T. Wada, A. S. Slutsky, D. Liu, C. Qin, C. Jiang and J. M. Penninger (2005). “A crucial role of angiotensin converting enzyme 2 (ACE2) in SARS coronavirus-induced lung injury.” Nat Med 11(8): 875–879.

Lan, J., J. Ge, J. Yu, S. Shan, H. Zhou, S. Fan, Q. Zhang, X. Shi, Q. Wang, L. Zhang and X. Wang (2020). “Structure of the SARS-CoV-2 spike receptor-binding domain bound to the ACE2 receptor.” Nature 581(7807): 215–220.

Lapish, C. C., S. Kroener, D. Durstewitz, A. Lavin and J. K. Seamans (2007). “The ability of the mesocortical dopamine system to operate in distinct temporal modes.” Psychopharmacology (Berl) 191(3): 609–625.

Leta, V., M. Rodriguez-Violante, A. Abundes, K. Rukavina, J. T. Teo, C. Falup-Pecurariu, L. Irincu, S. Rota, R. Bhidayasiri, A. Storch, P. Odin, A. Antonini and K. Ray Chaudhuri (2021). “Parkinson’s Disease and Post-COVID-19 Syndrome: The Parkinson’s Long-COVID Spectrum.” Mov Disord.

Li, W., M. J. Moore, N. Vasilieva, J. Sui, S. K. Wong, M. A. Berne, M. Somasundaran, J. L. Sullivan, K. Luzuriaga, T. C. Greenough, H. Choe and M. Farzan (2003). “Angiotensin-converting enzyme 2 is a functional receptor for the SARS coronavirus.” Nature 426(6965): 450–454.

Li, Z., J. Zhou, L. Lan, S. Cheng, R. Sun, Q. Gong, M. Wintermark, F. Zeng and F. Liang (2020). “Concurrent brain structural and functional alterations in patients with migraine without aura: an fMRI study.” J Headache Pain 21(1): 141.

Lu, R., X. Zhao, J. Li, P. Niu, B. Yang, H. Wu, W. Wang, H. Song, B. Huang, N. Zhu, Y. Bi, X. Ma, F. Zhan, L. Wang, T. Hu, H. Zhou, Z. Hu, W. Zhou, L. Zhao, J. Chen, Y. Meng, J. Wang, Y. Lin, J. Yuan, Z. Xie, J. Ma, W. J. Liu, D. Wang, W. Xu, E. C. Holmes, G. F. Gao, G. Wu, W. Chen, W. Shi and W. Tan (2020). “Genomic characterisation and epidemiology of 2019 novel coronavirus: implications for virus origins and receptor binding.” The Lancet 395(10224): 565–574.

Lukiw, W. J., A. Pogue and J. M. Hill (2020). “SARS-CoV-2 Infectivity and Neurological Targets in the Brain.” Cell Mol Neurobiol.

Mao, L., H. Jin, M. Wang, Y. Hu, S. Chen, Q. He, J. Chang, C. Hong, Y. Zhou, D. Wang, X. Miao, Y. Li and B. Hu (2020). “Neurologic Manifestations of Hospitalized Patients With Coronavirus Disease 2019 in Wuhan, China.” JAMA Neurol 77(6): 683–690.

Marshall, M. (2021). “COVID’s toll on smell and taste: what scientists do and don’t know.” Nature 589(7842): 342–343.

McKinley, M. J., A. L. Albiston, A. M. Allen, M. L. Mathai, C. N. May, R. M. McAllen, B. J. Oldfield, F. A. Mendelsohn and S. Y. Chai (2003). “The brain renin-angiotensin system: location and physiological roles.” Int J Biochem Cell Biol 35(6): 901–918.

Morassi, M., F. Palmerini, S. Nici, E. Magni, G. Savelli, U. P. Guerra, M. Chieregato, S. Morbelli and A. Vogrig (2021). “SARS-CoV-2-related encephalitis with prominent parkinsonism: clinical and FDG-PET correlates in two patients.” J Neurol.

Muus, C., M. D. Luecken, G. Eraslan, L. Sikkema, A. Waghray, G. Heimberg, Y. Kobayashi, E. D. Vaishnav, A. Subramanian, C. Smillie, K. A. Jagadeesh, E. T. Duong, E. Fiskin, E. T. Triglia, M. Ansari, P. Cai, B. Lin, J. Buchanan, S. Chen, J. Shu, A. L. Haber, H. Chung, D. T. Montoro, T. Adams, H. Aliee, S. J. Allon, Z. Andrusivova, I. Angelidis, O. Ashenberg, K. Bassler, C. Becavin, I. Benhar, J. Bergenstrahle, L. Bergenstrahle, L. Bolt, E. Braun, L. T. Bui, S. Callori, M. Chaffin, E. Chichelnitskiy, J. Chiou, T. M. Conlon, M. S. Cuoco, A. S. E. Cuomo, M. Deprez, G. Duclos, D. Fine, D. S. Fischer, S. Ghazanfar, A. Gillich, B. Giotti, J. Gould, M. Guo, A. J. Gutierrez, A. C. Habermann, T. Harvey, P. He, X. Hou, L. Hu, Y. Hu, A. Jaiswal, L. Ji, P. Jiang, T. S. Kapellos, C. S. Kuo, L. Larsson, M. A. Leney-Greene, K. Lim, M. Litvinukova, L. S. Ludwig, S. Lukassen, W. Luo, H. Maatz, E. Madissoon, L. Mamanova, K. Manakongtreecheep, S. Leroy, C. H. Mayr, I. M. Mbano, A. M. McAdams, A. N. Nabhan, S. K. Nyquist, L. Penland, O. B. Poirion, S. Poli, C. Qi, R. Queen, D. Reichart, I. Rosas, J. C. Schupp, C. V. Shea, X. Shi, R. Sinha, R. V. Sit, K. Slowikowski, M. Slyper, N. P. Smith, A. Sountoulidis, M. Strunz, T. B. Sullivan, D. Sun, C. Talavera-Lopez, P. Tan, J. Tantivit, K. J. Travaglini, N. R. Tucker, K. A. Vernon, M. H. Wadsworth, J. Waldman, X. Wang, K. Xu, W. Yan, W. Zhao, C. G. K. Ziegler, N. L. Consortium and N. Human Cell Atlas Lung Biological (2021). “Single-cell meta-analysis of SARS-CoV-2 entry genes across tissues and demographics.” Nat Med 27(3): 546–559.

Nagu, P., A. Parashar, T. Behl and V. Mehta (2021). “CNS implications of COVID-19: a comprehensive review.” Rev Neurosci 32(2): 219–234.

Nampoothiri, S., F. Sauve, G. Ternier, D. Fernandois, C. Coelho, M. Imbernon, E. Deligia, R. Perbet, V. Florent, M. Baroncini, F. Pasquier, F. Trottein, C.-A. Maurage, V. Mattot, P. Giacobini, S. Rasika and V. Prevot (2020). “The hypothalamus as a hub for putative SARS-CoV-2 brain infection.” bioRxiv: 2020.2006.2008.139329.

Onimaru, H., K. Ikeda, T. Mariho and K. Kawakami (2014). “Cytoarchitecture and CO(2) sensitivity of Phox2b-positive Parafacial neurons in the newborn rat medulla.” Prog Brain Res 209: 57–71.

Peiro, C. and S. Moncada (2020). “Substituting Angiotensin-(1-7) to Prevent Lung Damage in SARS-CoV-2 Infection?” Circulation 141(21): 1665–1666.

Pignatelli, A. and O. Belluzzi (2017). “Dopaminergic Neurones in the Main Olfactory Bulb: An Overview from an Electrophysiological Perspective.” Front Neuroanat 11: 7.

Prensa, L., J. M. Gimenez-Amaya, A. Parent, J. Bernacer and C. Cebrian (2009). “The nigrostriatal pathway: axonal collateralization and compartmental specificity.” J Neural Transm Suppl(73): 49–58.

Qiao, J., W. Li, J. Bao, Q. Peng, D. Wen, J. Wang and B. Sun (2020). “The expression of SARS-CoV-2 receptor ACE2 and CD147, and protease TMPRSS2 in human and mouse brain cells and mouse brain tissues.” Biochem Biophys Res Commun 533(4): 867–871.

Qin, Y., J. Wu, T. Chen, J. Li, G. Zhang, D. Wu, Y. Zhou, N. Zheng, A. Cai, Q. Ning, A. Manyande, F. Xu, J. Wang and W. Zhu (2021). “Long-term micro-structure and cerebral blood flow changes in patients recovered from COVID-19 without neurological manifestations.” J Clin Invest.

Rabelo, L. A., N. Alenina and M. Bader (2011). “ACE2-angiotensin-(1-7)-Mas axis and oxidative stress in cardiovascular disease.” Hypertens Res 34(2): 154–160.

Rahman, M. A., K. Islam, S. Rahman and M. Alamin (2021). “Neurobiochemical Cross-talk Between COVID-19 and Alzheimer’s Disease.” Mol Neurobiol 58(3): 1017–1023.

Reese, T. S. and M. J. Karnovsky (1967). “Fine structural localization of a blood-brain barrier to exogenous peroxidase.” J Cell Biol 34(1): 207–217.

Rice, G. I., D. A. Thomas, P. J. Grant, A. J. Turner and N. M. Hooper (2004). “Evaluation of angiotensin-converting enzyme (ACE), its homologue ACE2 and neprilysin in angiotensin peptide metabolism.” Biochem J 383(Pt 1): 45–51.

Sandgren, J. A., D. W. Linggonegoro, S. Y. Zhang, S. A. Sapouckey, K. E. Claflin, N. A. Pearson, M. R. Leidinger, G. L. Pierce, M. K. Santillan, K. N. Gibson-Corley, C. D. Sigmund and J. L. Grobe (2018). “Angiotensin AT1A receptors expressed in vasopressin-producing cells of the supraoptic nucleus contribute to osmotic control of vasopressin.” Am J Physiol Regul Integr Comp Physiol 314(6): R770–R780.

Sashindranath, M. and H. H. Nandurkar (2021). “Endothelial Dysfunction in the Brain: Setting the Stage for Stroke and Other Cerebrovascular Complications of COVID-19.” Stroke: STROKEAHA120032711.

Sashindranath, M. and H. H. Nandurkar (2021). “Endothelial Dysfunction in the Brain: Setting the Stage for Stroke and Other Cerebrovascular Complications of COVID-19.” Stroke 52(5): 1895–1904.

Satarker, S. and M. Nampoothiri (2020). “Involvement of the nervous system in COVID-19: The bell should toll in the brain.” Life Sci 262: 118568.

Schultz, W. (2002). “Getting formal with dopamine and reward.” Neuron 36(2): 241–263.

Segarra, M., M. R. Aburto and A. Acker-Palmer (2021). “Blood-Brain Barrier Dynamics to Maintain Brain Homeostasis.” Trends Neurosci 44(5): 393–405.

Smith, A. D. and J. P. Bolam (1990). “The neural network of the basal ganglia as revealed by the study of synaptic connections of identified neurones.” Trends Neurosci 13(7): 259–265.

Smith, J. C., A. P. Abdala, I. A. Rybak and J. F. Paton (2009). “Structural and functional architecture of respiratory networks in the mammalian brainstem.” Philos Trans R Soc Lond B Biol Sci 364(1529): 2577–2587.

Song, E., C. Zhang, B. Israelow, A. Lu-Culligan, A. V. Prado, S. Skriabine, P. Lu, O. E. Weizman, F. Liu, Y. Dai, K. Szigeti-Buck, Y. Yasumoto, G. Wang, C. Castaldi, J. Heltke, E. Ng, J. Wheeler, M. M. Alfajaro, E. Levavasseur, B. Fontes, N. G. Ravindra, D. Van Dijk, S. Mane, M. Gunel, A. Ring, S. A. J. Kazmi, K. Zhang, C. B. Wilen, T. L. Horvath, I. Plu, S. Haik, J. L. Thomas, A. Louvi, S. F. Farhadian, A. Huttner, D. Seilhean, N. Renier, K. Bilguvar and A. Iwasaki (2021). “Neuroinvasion of SARS-CoV-2 in human and mouse brain.” J Exp Med 218(3).

Sonkaya, A. R., B. OztUrk and S. O. Karada (2020). “Cerebral hemodynamic alterations in patients with Covid-19.” Turk J Med Sci.

Stamatakis, A. M. and G. D. Stuber (2012). “Activation of lateral habenula inputs to the ventral midbrain promotes behavioral avoidance.” Nat Neurosci 15(8): 1105–1107.

Stefano, G. B., R. Ptacek, H. Ptackova, A. Martin and R. M. Kream (2021). “Selective Neuronal Mitochondrial Targeting in SARS-CoV-2 Infection Affects Cognitive Processes to Induce ‘Brain Fog’ and Results in Behavioral Changes that Favor Viral Survival.” Med Sci Monit 27: e930886.

Swart, R. M., E. J. Hoorn, M. G. Betjes and R. Zietse (2011). “Hyponatremia and inflammation: the emerging role of interleukin-6 in osmoregulation.” Nephron Physiol 118(2): 45–51.

Tanaka, S. (2006). “Dopaminergic control of working memory and its relevance to schizophrenia: a circuit dynamics perspective.” Neuroscience 139(1): 153–171.

Tipnis, S. R., N. M. Hooper, R. Hyde, E. Karran, G. Christie and A. J. Turner (2000). “A human homolog of angiotensin-converting enzyme. Cloning and functional expression as a captopril-insensitive carboxypeptidase.” J Biol Chem 275(43): 33238–33243.

U, R. A. and K. Verma (2020). “Happy Hypoxemia in COVID-19-A Neural Hypothesis.” ACS Chem Neurosci 11(13): 1865–1867.

Uhal, B. D., X. Li, A. Xue, X. Gao and A. Abdul-Hafez (2011). “Regulation of alveolar epithelial cell survival by the ACE-2/angiotensin 1-7/Mas axis.” Am J Physiol Lung Cell Mol Physiol 301(3): L269–274.

Underwood, M. D., A. A. Khaibulina, S. P. Ellis, A. Moran, P. M. Rice, J. J. Mann and V. Arango (1999). “Morphometry of the dorsal raphe nucleus serotonergic neurons in suicide victims.” Biol Psychiatry 46(4): 473–483.

Verdecchia, P., C. Cavallini, A. Spanevello and F. Angeli (2020). “The pivotal link between ACE2 deficiency and SARS-CoV-2 infection.” Eur J Intern Med 76: 14–20.

Verharen, J. P. H., J. W. de Jong, T. J. M. Roelofs, C. F. M. Huffels, R. van Zessen, M. C. M. Luijendijk, R. Hamelink, I. Willuhn, H. E. M. den Ouden, G. van der Plasse, R. A. H. Adan and L. Vanderschuren (2018). “A neuronal mechanism underlying decision-making deficits during hyperdopaminergic states.” Nat Commun 9(1): 731.

Vezzani, A. and B. Viviani (2015). “Neuromodulatory properties of inflammatory cytokines and their impact on neuronal excitability.” Neuropharmacology 96(Pt A): 70–82.

Vickers, C., P. Hales, V. Kaushik, L. Dick, J. Gavin, J. Tang, K. Godbout, T. Parsons, E. Baronas, F. Hsieh, S. Acton, M. Patane, A. Nichols and P. Tummino (2002). “Hydrolysis of biological peptides by human angiotensin-converting enzyme-related carboxypeptidase.” J Biol Chem 277(17): 14838–14843.

Winter, S. M., J. Fresemann, C. Schnell, Y. Oku, J. Hirrlinger and S. Hulsmann (2009). “Glycinergic interneurons are functionally integrated into the inspiratory network of mouse medullary slices.” Pflugers Arch 458(3): 459–469.

Wise, R. A. (1978). “Catecholamine theories of reward: a critical review.” Brain Res 152(2): 215–247.

Xia, H., T. M. de Queiroz, S. Sriramula, Y. Feng, T. Johnson, I. N. Mungrue and E. Lazartigues (2015). “Brain ACE2 overexpression reduces DOCA-salt hypertension independently of endoplasmic reticulum stress.” Am J Physiol Regul Integr Comp Physiol 308(5): R370–378.

Yan, R., Y. Zhang, Y. Li, L. Xia, Y. Guo and Q. Zhou (2020). “Structural basis for the recognition of SARS-CoV-2 by full-length human ACE2.” Science 367(6485): 1444–1448.

Yousaf, Z., S. D. Al-Shokri, H. Al-Soub and M. F. H. Mohamed (2020). “COVID-19-associated SIADH: a clue in the times of pandemic!” Am J Physiol Endocrinol Metab 318(6): E882–E885.

Zhang, H., J. M. Penninger, Y. Li, N. Zhong and A. S. Slutsky (2020). “Angiotensin-converting enzyme 2 (ACE2) as a SARS-CoV-2 receptor: molecular mechanisms and potential therapeutic target.” Intensive Care Med 46(4): 586–590.

Zhang, L. and V. S. Hernandez (2013). “Synaptic innervation to rat hippocampus by vasopressin-immuno-positive fibres from the hypothalamic supraoptic and paraventricular nuclei.” Neuroscience 228: 139–162.

Zhang, L., V. S. Hernandez, E. Vazquez-Juarez, F. K. Chay and R. A. Barrio (2016). “Thirst Is Associated with Suppression of Habenula Output and Active Stress Coping: Is there a Role for a Non-canonical Vasopressin-Glutamate Pathway?” Front Neural Circuits 10: 13.

Zhang, L., M. A. Zetter, E. C. Guerra, V. S. Hernandez, S. K. Mahata and L. E. Eiden (2021). “ACE2 in the second act of COVID-19 syndrome: Peptide dysregulation and possible correction with oestrogen.” J Neuroendocrinol 33(2): e12935.

Zhao, H., B. L. Zhang, S. J. Yang and B. Rusak (2015). “The role of lateral habenula-dorsal raphe nucleus circuits in higher brain functions and psychiatric illness.” Behav Brain Res 277: 89–98.

